## Supplementary Information for "KAP1 negatively regulates RNA polymerase II elongation kinetics to activate signal-induced transcription"

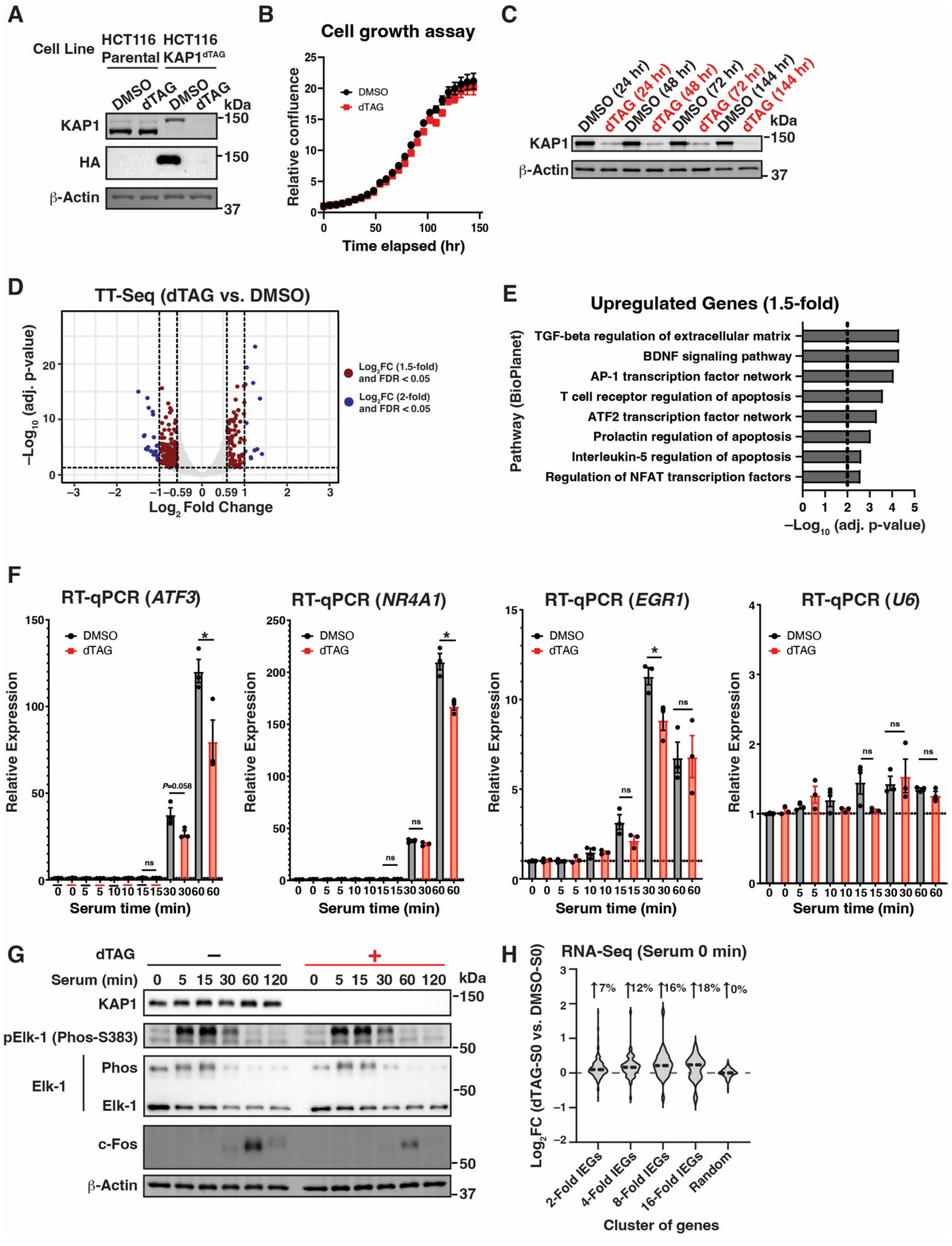


**Supplementary Figure 1. Acute KAP1 depletion leads to virtually no transcriptional changes in homeostatic conditions.** (**A**) dTAG characterization in HCT116 KAP1$^{dTAG}$

and HCT116 parental cells. (**B**) Cell growth assay showing relative cell confluence during a dTAG treatment time course. Data represent mean ± SEM (n=4). (**C**) Western blots to monitor KAP1 degradation kinetics up to 6 days of dTAG treatment. (**D**) TT-Seq volcano plot (n=2, false discovery rate [FDR] < 0.05). (**E**) Pathway analysis for 1.5-fold upregulated genes. (**F**) RT-qPCR assay highlighting gene expression of three representative IEGs (*ATF3, NR4A1, and EGR1*) and one control gene (*U6*) in DMSO- and dTAG-treated cells during a serum stimulation time course. Data represent mean ± SEM (n=3, Student's t-test comparing DMSO to dTAG at each of the indicated time points). **P*< 0.05, ***P*<0.01, ****P*<0.001, ns: not significant. (**G**) Western blots showing induction of phosphorylated Elk-1 upon serum stimulation time course with and without dTAG treatment. (**H**) Violin plot showing $Log_2FC$ values of DMSO versus dTAG treated cells before serum stimulation (serum 0 min). Median % changes in expression is listed above the violin in each gene cluster.

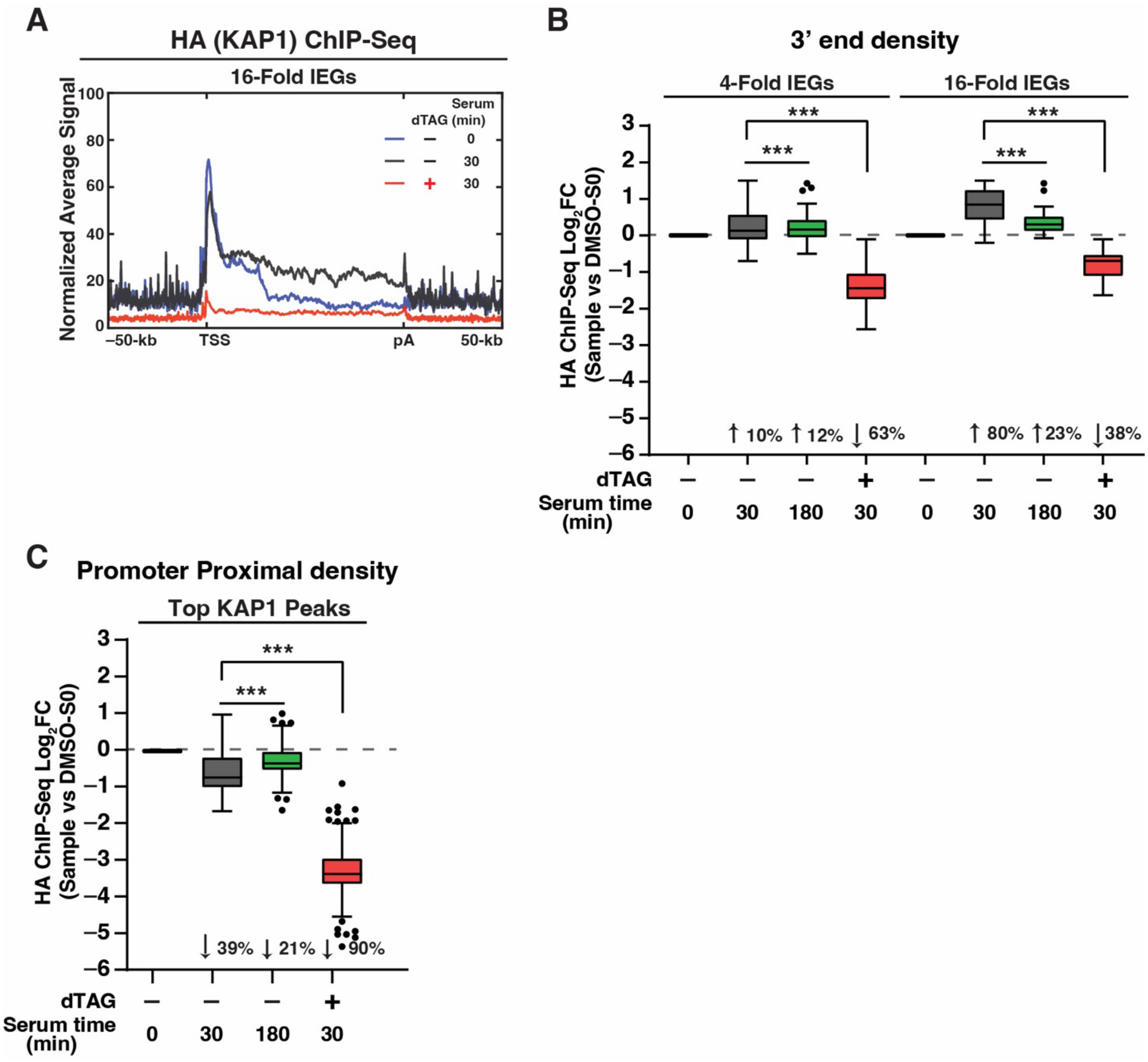


**Supplementary Figure 2. KAP1 localizes to the gene bodies and 3' ends of IEGs upon serum stimulation.** (**A**) HA (KAP1) ChIP-Seq metagene analysis at 16-Fold IEGs in the three indicated conditions with an extended X-axis to show 50-kb upstream the TSS and downstream the pA site. (**B**) HA ChIP-Seq quantitation's of KAP1 density at the 3' ends of IEGs (4-Fold and 16-Fold). The $Log_2FC$ value is plotted for the respective time point normalized to the DMSO-Serum 0 min (S0) sample. Statistical analysis was performed between the dTAG-Serum condition and the respective condition shown on the Tukey test plot. Wilcoxon signed-rank test (*$P$<0.05, **$P$<0.01, ***$P$<0.001). (**C**) HA ChIP-Seq quantitation's of KAP1 density at the top KAP1 peaks at PP regions (n=181, 19 genes were removed from the top 200 peaks because they contained no signal in at least one condition). The $Log_2FC$ value is plotted for the respective time point normalized to the DMSO-Serum 0 min (S0) sample. Statistical analysis was performed between the dTAG-Serum condition and the respective condition shown on the Tukey test plot. Wilcoxon signed-rank test (*$P$<0.05, **$P$<0.01, ***$P$<0.001).

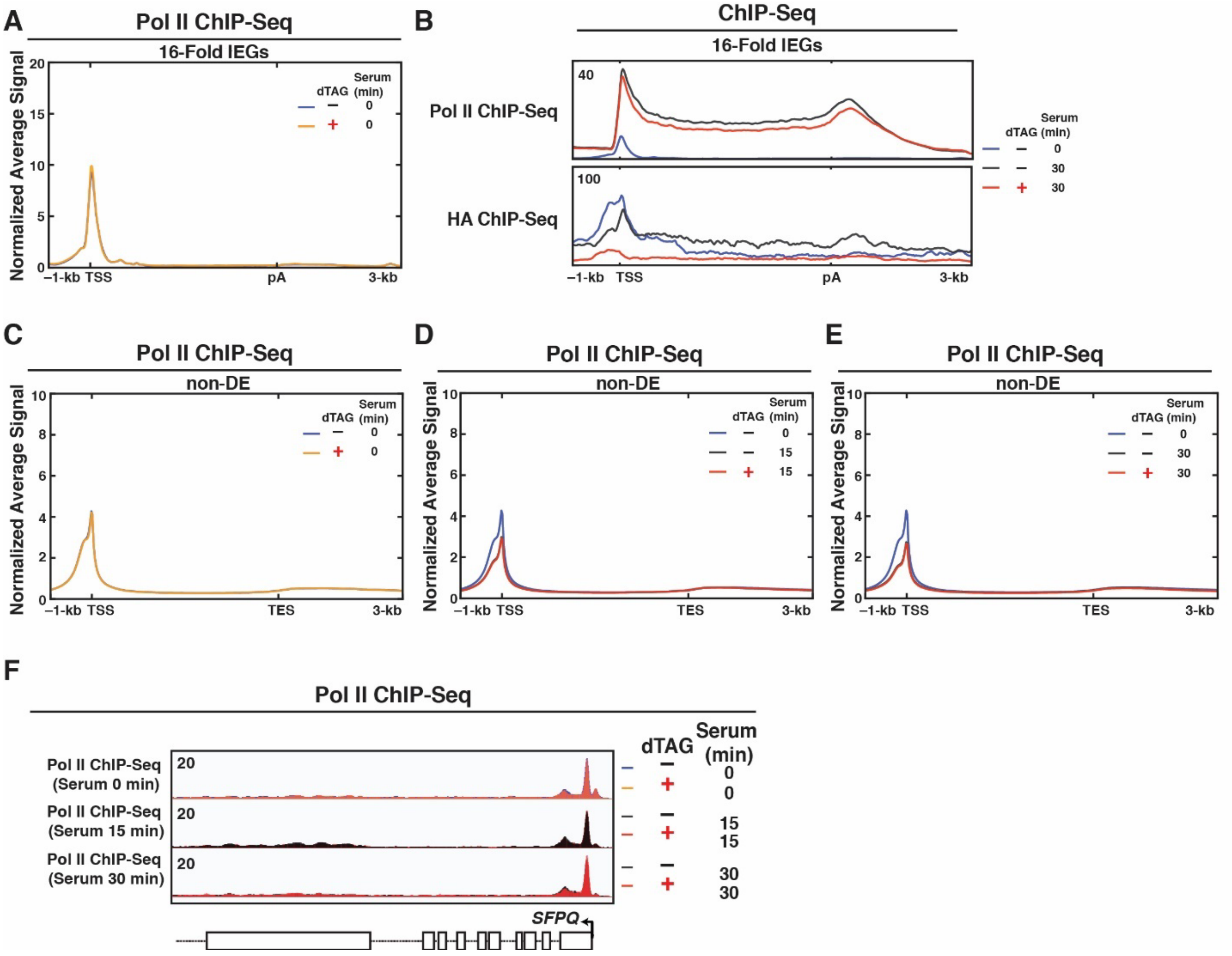


**Supplementary Figure 3. KAP1 regulates Pol II occupancy during serum stimulation.** (**A**) Pol II ChIP-Seq metagene analysis at 16-Fold IEGs in the serum 0 min condition with and without dTAG treatment. (**B**) Overlayed metagene profiles of Pol II and HA (KAP1) ChIP-Seq in the three indicated conditions. (**C-E**) Pol II ChIP-Seq metagene profiles at non-DE genes: (**C**) 0 min, (**D**) 15 min, (**E**) and 30 min serum stimulation time points. (**F**) Pol II ChIP-Seq browser tracks in multiple conditions at one control gene (*SFPQ*).

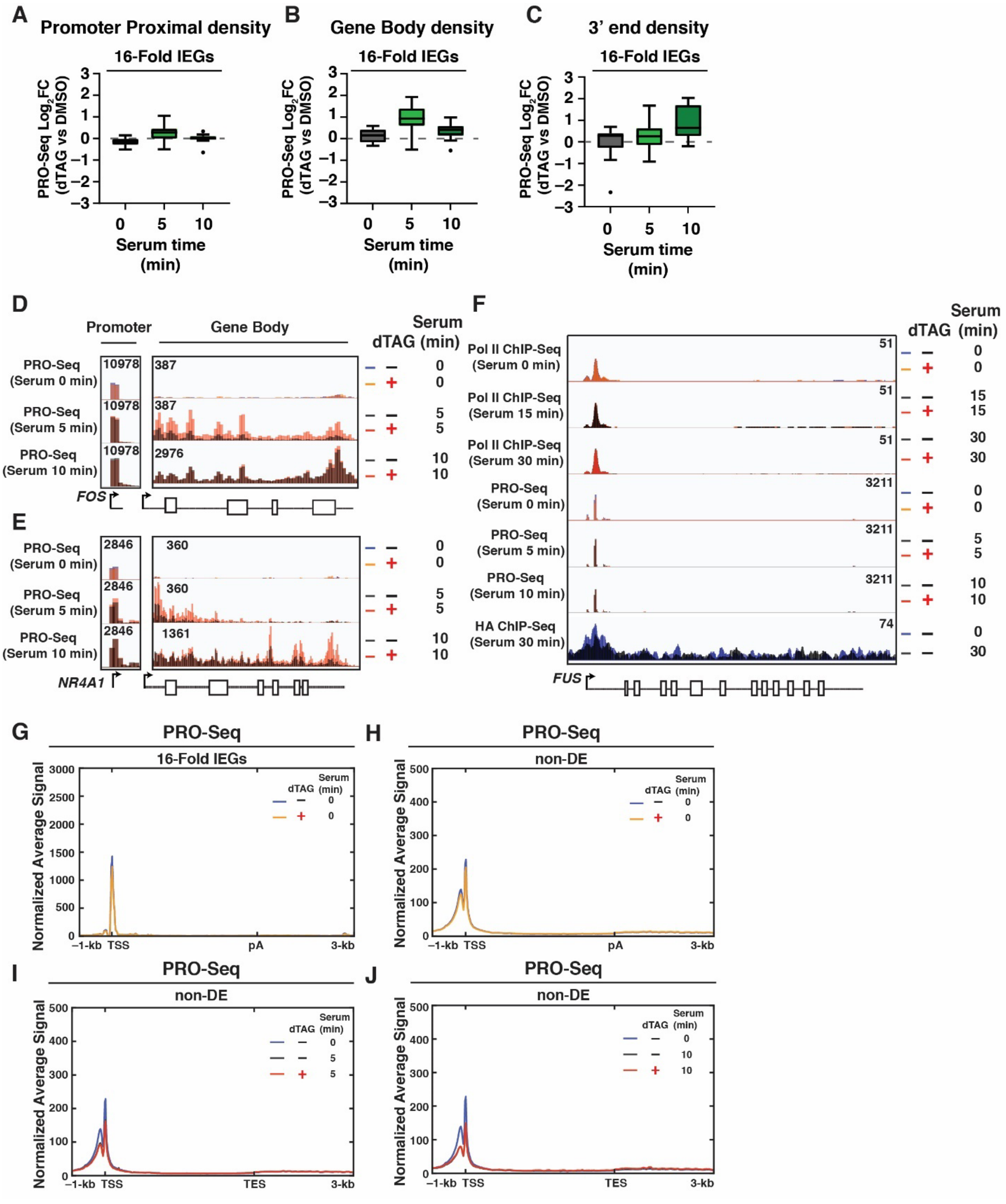


**Supplementary Figure 4. Acute KAP1 depletion leads to increased active Pol II density in the gene bodies of IEGs upon serum stimulation.** (**A-C**) Quantitation's of PRO-Seq signal at IEGs (16-Fold): (**A**) PP regions, (**B**) GB regions, and (**C**) 3' ends. The $Log_2FC$ value is plotted for dTAG versus DMSO at the respective serum time point. (**D-E**) PRO-Seq browser tracks at multiple time points at the (**D**) *FOS* locus and (**E**) *NR4A1*

locus. (**F**) Pol II ChIP-Seq, PRO-Seq, and HA ChIP-Seq browser tracks in multiple conditions at a control gene (*FUS*). (**G-H**) PRO-Seq metagene analysis at 16-Fold IEGs (**G**) and non-DE genes (**H**) in 0 min serum stimulation condition. (**I-J**) PRO-Seq metagene analysis at non-DE genes at the (**I**) 5 min and (**J**) 10 min serum stimulation time points.

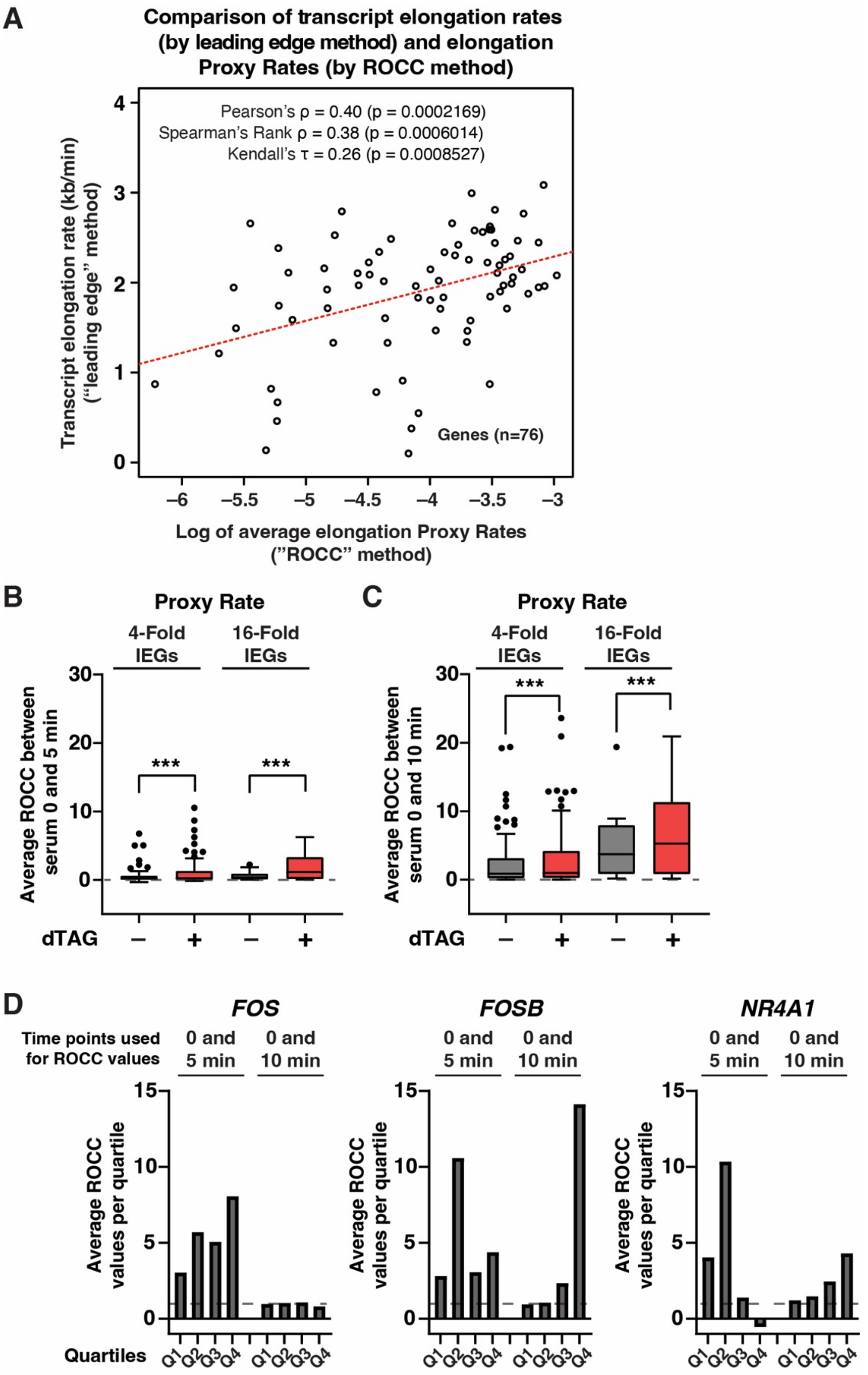


**Supplementary Figure 5. Benchmarking of the Proxy Rate analysis using ROCC.** (**A**) Validation of the ROCC approach to estimate elongation rate using a published estrogen-inducible gene dataset[1]. The ROCC Proxy Rate analysis was performed as described and tested for correlation to the leading edge method, which captures the wavefront to calculate the elongation rate. (**B-C**) Proxy Rate calculations between the (**B**) serum 0 and 5 min time points, and (**C**) serum 0 and 10 min time points, separately. This analysis simply subtracts signal between the designated time point thus allowing the serum 0 to exert maximal influence on the overall rate analysis. Wilcoxon signed-rank test (**P*<0.05, ***P*<0.01, ****P*<0.001). (**D**) Proxy Rate calculations for representative IEGs across the four quartiles of gene length. This analysis uses the "single time point" ROCC approach plotted in panels (**B**) and (**C**). The single time points are designated at the top of the graph. Q1: quartile 1, Q2: quartile 2, Q3: quartile 3, Q4: quartile 4.

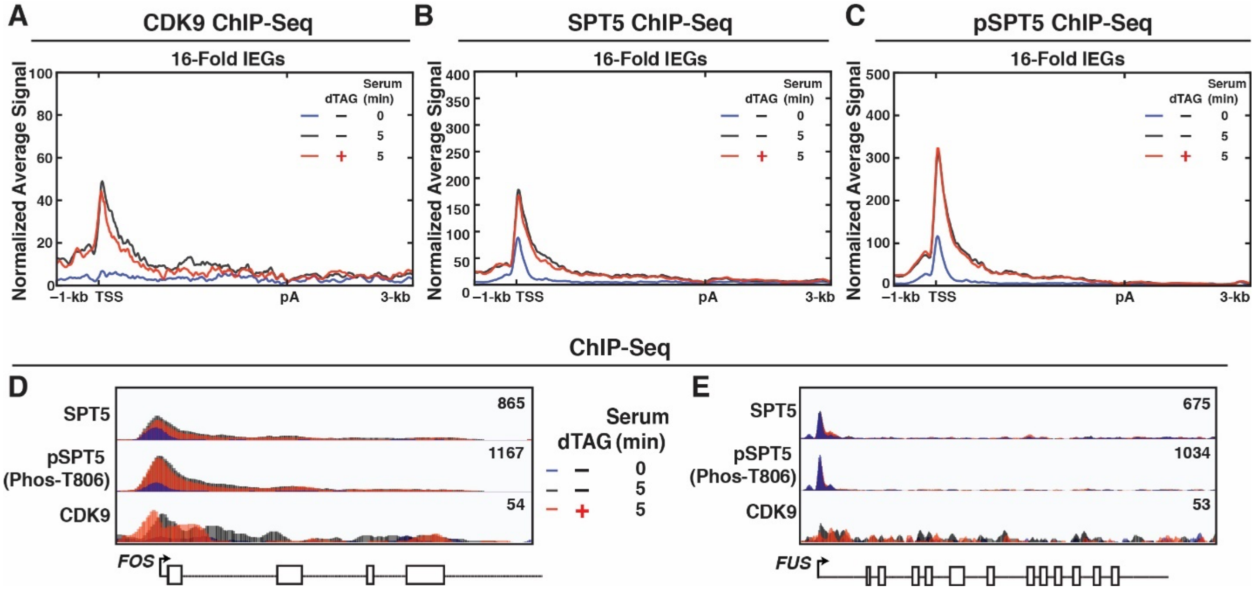


**Supplementary Figure 6. Acute KAP1 depletion does not affect the recruitment nor the phosphorylation status of pause release factors.** (**A-C**) ChIP-Seq metagene profile of (**A**) CDK9, (**B**) total SPT5, and (**C**) pSPT5 (Phos-T806) at 16-Fold IEGs in the three indicated conditions. (**D-E**) CDK9, SPT5, and pSPT5 browser tracks of (**D**) one representative IEG (*FOS*) and (**E**) one control gene (*FUS*).

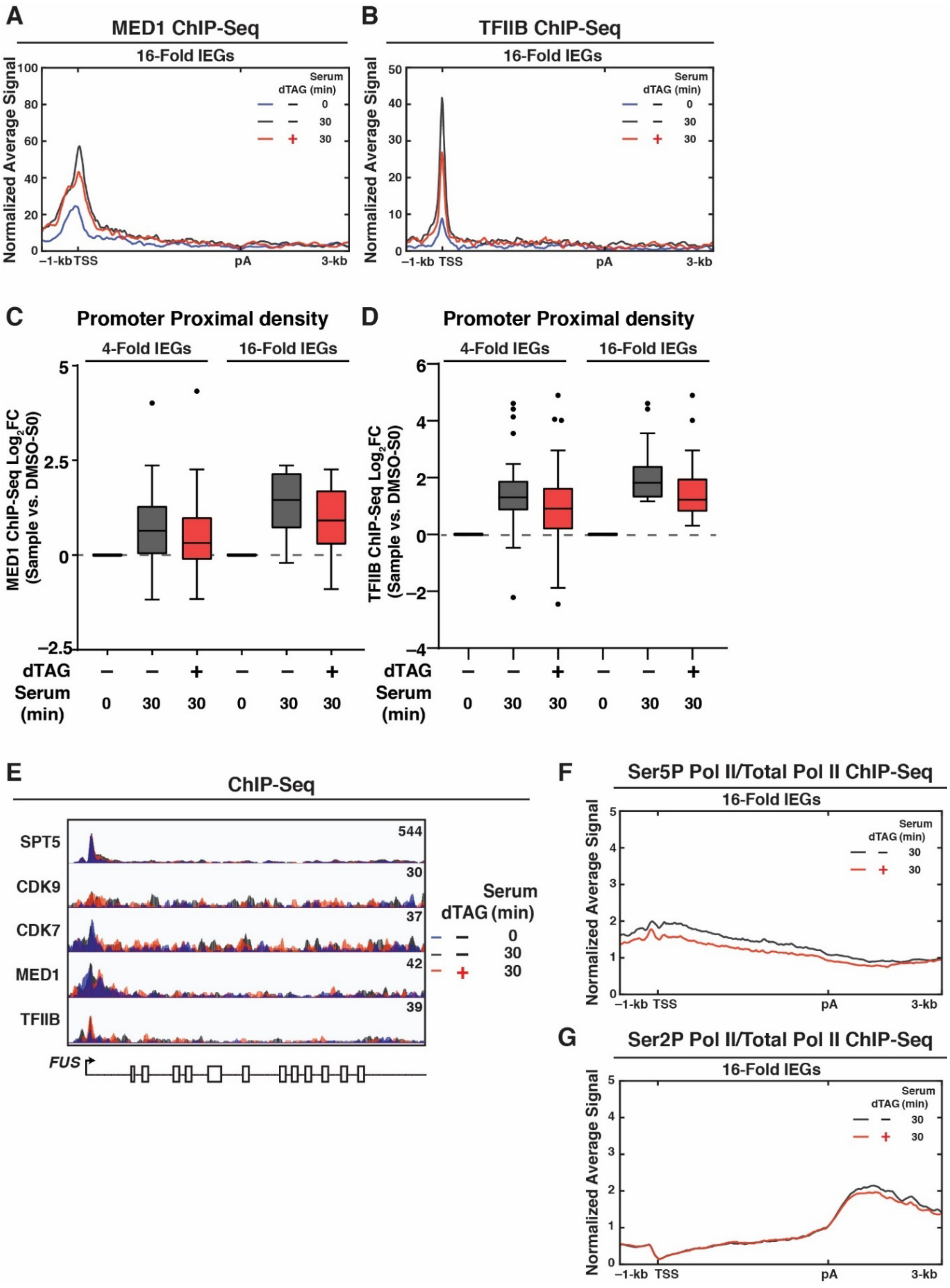


**Supplementary Figure 7. Acute KAP1 depletion leads to decreased occupancy of regulators of transcription elongation and initiation.** (**A-B**) ChIP-Seq metagene analysis of (**A**) MED1 and (**B**) TFIIB at 16-Fold IEGs in the three indicated conditions. (**C-D**) ChIP-Seq quantitation of (**C**) MED1 and (**D**) TFIIB in PP regions. $Log_2FC$ value is plotted to compare each time point to the DMSO-Serum 0 min sample. The median percent increase when compared to DMSO-Serum 0 is listed below each bar in the Tukey plot. (**E**) ChIP-Seq browser tracks of all factors in multiple conditions at one control gene (*FUS*). (**F-G**) ChIP-Seq metagene analysis of (**F**) Ser5P Pol II (normalized to total Pol II) and (**F**) Ser2P Pol II (normalized to total Pol II) at 16-Fold IEGs in DMSO-Serum 30 min and dTAG-Serum 30 min conditions.

### Supplementary Methods

**Cloning of dTAG constructs.** For cloning of dTAG constructs as well as making the HCT116 KAP1$^{dTAG}$ cell line, published procedures were followed with minor modifications[2]. Briefly, the gRNA was designed from a combination of Benchling for off-target scores and the Washington University gRNA designer tool for structural scores (https://crisprdb.org/wu-crispr-website/index.html). The primers for both gRNA and arm cloning were designed based on this gRNA selection following the tutorial[2]. To clone the gRNA plasmid, custom made DNA oligonucleotides (MilliporeSigma) were annealed with T4 ligase (NEB, catalog M0202S) and phosphorylated with T4 polynucleotide kinase (NEB, catalog M0201L) by incubating at 37ºC for 30 min, 95ºC for 5 min, and a ramp down to 25ºC at 5ºC/min. To prepare the sgRNA backbone vector, the empty universal cutting vector (pX330A-sgX-sgPITCh) was digested with BbsI-HF (NEB, catalog R0539S) for 1 hr at 37ºC followed by a 15 min treatment with Quick CIP (NEB, catalog M0525S) and subsequent gel purification (QIAGEN, catalog 28606). Prepared oligonucleotides were then ligated into the digested backbone vector with T4 ligase at room temperature for 1 hr followed by heat-inactivation and transformation into home-made DH5$\alpha$ competent cells followed by plasmid preparation and sequence verification using the primers listed in Supplemetary Table 2. To clone arm plasmids, the pCRIS-PITChv2-dTAG-Puro and pCRIS-PITChv2-dTAG-BSD plasmids were digested with MluI-HF (NEB, R3198S) followed by gel purification to prepare the linearized vector. To prepare the arm cassette, the original arm plasmid was utilized as template for PCR using the designed arm primers to insert the KAP1 C-terminal homology arms using Q5® Hot Start High-Fidelity DNA Polymerase (NEB, catalog M0493L). The ~1-kb product was purified and Gibson assembly performed (50°C, 30 min) using 1.5 µL of NEBuilder HiFi Master Mix (NEB, catalog E2621S) and the digested vector and PCR product. The Gibson product was diluted and transformed into home-made DH5$\alpha$ competent cells, and plasmid DNAs were prepared and verified by Sanger sequencing. All plasmids and primers used in this study are listed in Supplementary Table 1 and 2, respectively.

**Creation of HCT116 KAP1$^{dTAG}$ cells.** To generate HCT116 KAP1$^{dTAG}$ cells, we used a double selection approach in which both puromycin- and blasticidin-resistant dTAG cassette plasmids were transfected in tandem with a sgRNA plasmid. We selected cells with both antibiotics to target the two *KAP1/TRIM28* alleles and single-sorted for expansion followed by western blot to ensure no presence of the endogenous (lower molecular weight) KAP1 protein band. Briefly, HCT116 parental cells were seeded in 6-well plates (~500,000 cells/well) and transfected ~24 hr post-seeding with three plasmids (BSD arm plasmid, Puro arm plasmid, and gRNA plasmid) targeting the C-terminus of KAP1 at a ratio of 1:1:1 (0.33 µg each). Before transfection, the media was changed into 1 mL of fresh DMEM. Two mixes were prepared: (1) a DNA mix consisting of 1 µg of total plasmid DNA (0.33 µg of each plasmid) into 100 µL of OPTIMEM (Gibco, catalog 11058021) and a Lipofectamine 2000 (ThermoFisher Scientific, catalog 52887) mix of 3 µL incubated in final 100 µL OPTIMEM. The Lipofectamine 2000 mix was added dropwise to the DNA mix and incubated for 10 min at room temperature. After incubation, the 200 µL mix was added dropwise to cells and media was changed into 2 mL of DMEM media after 5 hr. Two days post-transfection, selection media (1.5 µg/mL puromycin

(ThermoFisher Scientific, catalog 227420500) and 10 µg/mL blasticidin (ThermoFisher Scientific, catalog BP264750) in complete DMEM) was added to the cells for 14 days to induce targeting of both *KAP1*/*TRIM28 alleles*, at which point stable colonies began to expand. A single-selection approach including 0.5 µg of only PURO arm plasmid and 0.5 µg of gRNA plasmid was included as control. Cells were re-split in a 6-well plate, grown for 4 days, and then expanded to a 10-cm tissue culture dish. Population lysates were collected and subjected to KAP1 western blot to verify targeting efficiency. Cells were single-cell sorted in 96-well plates at the UTSW Childrens Research Institute Moody Foundation Flow Cytometry Core in complete DMEM (200 µL per well with no antibiotic). Cells grew in single-cell format for ~4-5 weeks prior to slowly expanding to 10-cm tissue culture dishes (24-well first, 6-well second, and then finally to 10-cm dishes). To verify *KAP1* targeting, anti-KAP1 westerns blots were performed.

**Cell Growth assay.** For the cell growth assay, cells were treated with either 8 hr DMSO or dTAG and then the IncuCyte S3 Live Cell Imaging System (Essen Bioscience) was used for cell proliferation assays. 2000 cells/well were seeded into 96-well plates and imaged every 6 hr over 144 hr (6 days). Phase contrast images were used to calculate cell confluence using the IncuCyte software and the cell confluence at each time point was normalized to time 0.

**RNA extraction and RT-qPCR.** For RT-qPCR, cells were treated as described and RNA was extracted using the Zymo Quick-RNA MiniPrep Kit (Zymo Research, catalog R1055) following the kit instructions. RNA was quantified using the DeNovix DS-II FX+ Spectrophotometer and all RNAs were diluted to the same yield and re-quantified prior to cDNA synthesis reaction. To prepare cDNA, 2 µg of RNA was incubated with 1/200$^{th}$ of a unit of hexanucleotide primers (MilliporeSigma, catalog H0268-1UN) and 1 µL of 10 mM dNTP mix (NEB, catalog N0447L) for 5 min at 70°C. Next, 2 µL of 10X M-MuLV Reverse Transcriptase buffer and 1 µL of M-MuLV Reverse Transcriptase (NEB, catalog M0253L) was added to each sample (final volume 20 µL) and incubated at 42°C for 1 hr. The reaction was inactivated at 70°C for 10 min and samples were diluted by adding 80 uL with $H_2O$. For qPCR, 1 µL of diluted cDNA, 5 µL PowerUp™ SYBR™ Green Master Mix for qPCR (ThermoFisher Scientific, catalog A25741), 3 µL of $H_2O$, and 1 µL of 5 µM primer mix was used for each well in a 96-well plate. All primers used for RT-qPCR analysis are listed in Supplemetary Table 2. Samples were amplified for 40 cycles using the Applied Biosystems QuantStudio™ 3 Real-Time PCR System. All RT-qPCR data was analyzed using the ΔΔCt method where each RNA for each sample was normalized to the DMSO-Serum 0 min (S0) sample and then each target gene was normalized to two control genes (*RPL19* and *GAPDH*) using the geometric mean of their expression.

**RNA-Seq.** RNAs for RNA-Seq were prepared as described for RT-qPCR. For RNA-Seq library preparation, 1 µg of RNA was used as input with 2 µL of a 1:100 dilution of ERCC RNA Spike-In Mix (ThermoFisher Scientific, catalog 4456740) and the KAPA RNA Hyper+RiboErase HMR kit was used (Roche, catalog 8098131702) following the manufacturer's instructions for library preparation using KAPA beads (Roche, catalog KK8001). The following steps were performed: Oligo hybridization and rRNA depletion, KAPA bead cleanup, digestion with DNAse, and another KAPA bead cleanup as described in the kit protocol. RNA elution, fragmentation (8 min at 94°C) and priming was performed followed by first-strand synthesis and second-strand synthesis/A-tailing. Adapter ligation (15 min at 20°C) was performed with 7 µM NEBNext® Adaptor for Illumina® (NEB, kit catalog E6446L) followed by a 3 µL USER enzyme digest (15 min at 37°C). This was immediately followed by two cleanups (first with 0.63X KAPA beads and second with 0.7X PEG/NaCl solution). After adapter ligation, samples were PCR amplified using Illumina indexed primers (NEB, E7335L) for 8 cycles. Following a final KAPA bead purification and elution with 20 µL of 10 mM Tris-HCl pH = 8.0, size distribution and quality of libraries was determined using the Agilent Tapestation DNA ScreenTape (Agilent, catalog 5067-5582). Libraries were then quantified using the Qubit dsDNA HS Assay (ThermoFisher Scientific, catalog Q32851) and sequenced at ~33E6 paired-end reads/sample with 50-bp length using a NextSeq 500 (Illumina) at the UT Southwestern McDermott Center Sequencing Core. Three biological replicates per treatment condition were submitted.

**TT-Seq.** For TT-Seq, the protocol from Patrick Cramer's lab was followed with minor modifications[3]. Three 10-cm tissue culture dishes per condition were seeded such that cells reached 75% confluence 3 days post-seeding. On the third day, cells were treated with DMSO or dTAG for 8 hr, after which 500 µM 4-Thiouridine (MilliporeSigma, catalog T4509) was added to the media for 10 min. An extra dish was seeded per condition and ~12E6 cells were counted per plate and thus 36E6 cells/sample were used to prepare RNA. Cells were washed with 5 mL of 1X PBS once and then immediately 1 mL of TRIzol™ Reagent (ThermoFisher Scientific, catalog 15596026) was added and followed by 10 min of rocking. Cells were scaped off the plate, transferred to a 15 mL tube, and then homogenized by pipetting ~10 times. Cells were placed in –80°C until ready for RNA extraction. To isolate total RNA, 200 µL chloroform was added to the 1 mL of sample (3 tubes per replicate) and vortexed 15 sec followed by centrifugation (13,000xg, 15 min, 4°C). The upper, aqueous layer (~490 uL) was added to a fresh tube with 1 µL Glycoblue and an equal volume of Isopropanol (~490 µL) followed by mixing, incubation at room temperature for 10 min, and then centrifugation (13,000xg, 10 min, 4°C). The pellet was washed twice with 1 mL 75% Ethanol followed by a centrifugation (7500xg, 5 min, 4°C). After drying, the pellet was resuspended in 320 µL RNAse-free water per replicate, quantified by nanodrop, and diluted to 750 ng/µL followed by sonication using a Q800R3 Qsonica (50% amplitude with 3 cycles 30 seconds ON, 30 seconds OFF at 4°C). The RNA was denatured at 65°C for 10 min, placed on ice for 5 min, and diluted to 150 µg/700 µL to prepare for biotinylation. 10X Biotinylation buffer (100 mM Tris-HCl pH = 7.5, 10 mM EDTA pH = 8.0) was prepared along with 5X (1 mg/mL) EZ-Link™ HPDP-Biotin (MilliporeSigma, catalog 21341) in N,N-Dimethylformamide (Acros Organics, catalog 279600010). 300 µgs total RNA was used per replicate, and thus each condition had 2 x 1000 µL biotinylation reactions. A mix of Biotinylation Buffer (100 µL per reaction) and Biotin-HPDP was made (200 µL per reaction) and added to 700 µL of diluted RNA, the samples were mixed and then nutated for 2 hr at room temperature while covered in foil. To purify RNA after biotinylation, 500 µL of each reaction was transferred to a new tube followed by addition of equal volume of chloroform (500 µL). Samples were vortexed, incubated for 3 min, and then centrifuged (1,500xg, 5 min, 4°C), followed by transfer of the aqueous layer (~300 µL) to 2 new tubes per sample (~600 µL each) and then precipitation with 1/10$^{th}$ volume 5 M NaCl (~60 µL), 1 µL glycoblue, and 1X isopropanol

(~600 µL). Samples were vortexed, centrifuged (13,000xg, 30 min, 4°C), washed two times with 75% ethanol, and then resuspended in 500 µL of RPB buffer (10 mM Tris-HCl pH = 7.5, 1 mM EDTA pH = 8.0, 300 mM NaCl). To isolate biotinylated RNA, 200 uL of Dynabeads MyOne Streptavidin T1 (ThermoFisher Scientific, catalog 65601) was aliquoted per sample and washed 4 times with 1 mL beads wash buffer (10 mM Tris-HCl pH = 7.5, 1 mM EDTA pH = 8.0, 50 mM NaCl) and finally suspended in 1 mL beads wash buffer per sample plus 0.1% polyvinylpyrrolidone (ThermoFisher Scientific, catalog BP431-100). Beads were then nutated at room temperature for 10 min, washed one time with 1 mL beads wash buffer, and then suspended in RPB (200 µL per reaction). The biotinylated RNA product was denatured at 65°C for 5 min, placed on ice for 2 min, and then 200 µL of blocked beads to each sample was added followed by a 30 min incubation with rotation at room temperature. After 30 min, beads were washed 5 times with 4sU wash buffer (10 mM Tris-HCl pH = 7.5, 1 mM EDTA pH = 8.0, 1 M NaCl, 0.1% Tween-20). Biotinylated, 4sU-RNA was eluted from beads twice by adding 75 µL of 0.1 M DTT to beads and nutating for 15 min at room temperature, yielding a final elution volume of ~150 µL. 4sU-RNA was purified and concentrated using the Zymo RNA Clean and Concentrator Kit (Zymo Research, catalog R1013) following kit instructions and RNA was quantified using the Qubit™ RNA High Sensitivity (HS) Assay (ThermoFisher Scientific, Q32852). Libraries were prepared similarly to RNA-Seq except with the minor modifications described here. First, 2 µL of a 1:4000 ERCC Spike-in mix was added to approximately ~50 ngs of nascent RNA as input to library preparation. Fragmentation was done for 6 min at 94°C and 13 cycles of PCR were used for PCR amplification step. Libraries were quality checked with Tapestation and Qubit and then sequenced at ~100E6 paired-end reads/sample with 50-bp length using a NextSeq 500 (Illumina) at the UT Southwestern McDermott Center Sequencing Core. Two biological replicates per treatment condition were submitted.

**ChIP-Seq.** All ChIP-Seq experiments were performed as previously described[4] with minor modifications. For each ChIP, 1 x 15-cm tissue culture dish (yielding ~30E6 cells/plate) was utilized and seeded according to the experimental design in each figure (DMSO/dTAG +/- serum at different time points), and 20E6 cells were utilized per ChIP. For all Pol II ChIPs (RPB3, Ser5P Pol II, Ser2P Pol II), cells were crosslinked with 0.5% methanol-free formaldehyde (ThermoFisher Scientific, 28908) by adding directly to the tissue culture dish in media at room temperature for 10 min with rocking and neutralization with 150 mM glycine for 5 min with rocking. For all other ChIP-Seq experiments (HA (KAP1), SPT5, pSPT5, CDK9, CDK7, TFIIB, MED1), 1% formaldehyde for 10 min was used. For all experiments, the media was removed and cells were washed twice with cold 1X PBS, cold PBS was then added to the plate and then cells were collected by scraping. Cells were centrifuged (1000xg, 5 min, 4°C), and pelleted, flash frozen in liquid nitrogen, and frozen at –80°C until ready to use. To perform the ChIP, cells were resuspended in 4 mL/dish of Farnham Lysis Buffer (5 mM PIPES pH = 8.0, 85 mM KCl, 0.5% NP-40, 1 mM PMSF, 1X Protease Inhibitor (RPI, catalog P50900-1)), counted by hemocytometer, resuspended to 10E6 cells/mL, nutated for 30 min at 4°C, and then centrifuged to isolate nuclei (1000xg, 5 min, 4°C). The supernatant was removed and nuclei resuspended in Szak's RIPA Buffer (50 mM Tris-HCl pH = 8.0, 1% NP-40, 150 mM NaCl, 0.5% Na-Deoxycholate, 0.1% SDS, 2.5 mM EDTA pH = 8.0, 1 mM PMSF, and 1X Protease

Inhibitor) at a concentration of 25E6 nuclei/mL. The chromatin was sheared using a Q800R3 Qsonica (50% amplitude with 25 cycles 30 seconds ON, 30 seconds OFF at 4°C) to a DNA molecular weight range of 200-400-bp. After sonication, chromatin was centrifuged (21,000xg, 15 min, 4°C) and the supernatant was taken as clarified chromatin. For ChIP-Seq experiments that contained Drosophila spike-in, 50 ng of spike-in chromatin (Active Motif, catalog 53083) was added to each chromatin sample. Sheared chromatin was pre-cleared by incubating with 25 µL of Szak's RIPA equilibrated Protein G Dynabeads (ThermoFisher Scientific, 10003D) for 1 hr at 4°C. To equilibrate beads for pre-clearing, a master mix of beads were incubated with RIPA buffer, nutated for 5 min, and placed on a magnet to remove supernatant three times. To bind antibody to beads, 100 µL of Protein G Dynabeads per sample were equilibrated with 1X PBS+0.05% Tween-20 and resuspended to a final volume of 250 µL per ChIP. The corresponding antibody (see Supplemetary Table 3 for antibody details) was then added to the 250 µL beads, and the bead-antibody mix was nutated for 1 hr at 4°C. For samples that contained Drosophila spike-in, 2.5 µg of antibody specific for Drosophila H2Av (Active Motif, catalog 61686) was added to each tube. Antibody bound beads were then blocked in Szak's RIPA Buffer + 5% BSA for 1 hr at 4°C with rotation. Pre-cleared sheared chromatin was then added to beads and incubated overnight at 4°C with rotation. Beads from each sample were washed 2 times with 900 µL of Szak's RIPA Buffer, Low Salt Buffer (0.1% SDS, 1% NP-40, 2 mM EDTA pH = 8.0, 20 mM Tris-HCl pH = 8.0, 150 mM NaCl), High Salt Buffer (0.1% SDS, 1% NP-40, 2 mM EDTA pH = 8.0, 20 mM Tris-HCl pH = 8.0, 500 mM NaCl), LiCl buffer (250 mM LiCl, 1% NP-40, 1% sodium deoxycholate, 1 mM EDTA pH = 8.0, 20 mM Tris-HCl pH = 8.0), and TE Buffer (10 mM Tris-HCl pH = 8.0, 1 mM EDTA pH = 8.0). After the final wash, samples were pulse spun in a table-top centrifuge to get rid of residual buffer before placing on the magnet. Samples were then eluted from beads in 100 µL of elution buffer (100 mM $NaHCO_3$ pH = 8.0, 1% SDS) for 30 min at 65°C while vortexing every ~10 min. Input samples (40 µL) were volumed up to 100 µL by adding 60 µL elution buffer. DNA was eluted by placing the beads on a magnet, elutions transferred to new tubes, and de-crosslinked for 4 hr at 65°C with 100 µL volume of de-crosslinking buffer (500 mM NaCl, 2 mM EDTA pH = 8.0, 20 mM Tris-HCl pH = 6.8, 0.5 mg/mL Proteinase K (Epicentre, catalog MPR-90938)). ChIP DNA was purified and concentrated with the Zymo ChIP DNA Clean & Concentrator (Zymo Research, catalog D5201). ChIP samples were quantified by Qubit, and libraries were prepared using the KAPA Hyper Prep Kit (KAPA Biosystems, catalog KK8502) following the manufacturer's instructions. Briefly, samples underwent End Repair & A-tailing followed by adapter ligation with 300 nM to 1.5 µM NEB adapter depending on initial yield. Following a post-ligation cleanup, PCR amplification was performed with anywhere between 8-14 total cycles depending on initial yield. A KAPA bead cleanup was performed (1X) followed by size selection. For size selection, the first cleanup utilized 35 µL KAPA beads where the final supernatant contains the DNA of interest. The second and final cleanup utilized 10 µL of beads where the beads contain the bound, desired DNA. Following elution with 20 µL of 10 mM Tris-HCl pH = 8.0, libraries were quality-checked with Tapestation and Qubit, and then sequenced at ~33E6 paired-end reads/sample with 50-bp length using a NextSeq 500 (Illumina) at the UT Southwestern McDermott Center Sequencing Core. Two biological replicates of each ChIP-Seq for HA, Pol II, and Ser2P Pol II were completed, while one replicate of SPT5, pSPT5, CDK9, CDK7, TFIIB, MED1, and Ser5P Pol II were performed in each condition.

**PRO-Seq.** For PRO-Seq experiments, we followed the qPRO-Seq protocol[5] with minor modifications. Cells were seeded in 10-cm tissue culture dishes and treated as indicated and were ~90% confluent prior to collection. For each PRO-Seq, 4E6 cells were utilized. To prepare permeabilized cells for run-on, cells were first washed twice on the plate with 5 mL of ice-cold 1X PBS. Then, 2.5 mL of ice-cold Cell Permeabilization Buffer (CPB) (10 mM Tris-HCl pH = 8.0, 250 mM Sucrose, 10 mM KCl, 5 mM $MgCl_2$, 1 mM EGTA pH = 8.0, 0.1% NP-40, 0.5 mM DTT, 0.05% Tween-20, 0.1% Triton X-100, 10% Glycerol, 1X protease inhibitor, and 2 µL SUPERase-In RNase inhibitor (ThermoFisher Scientific, catalog AM2696) per 10 mL) was immediately added to the tissue culture dish. Triton X-100 was added to CPB to increase permeabilization. The tissue culture dishes were placed on ice and then immediately scraped with a cell scraper. At this point, cells were collected in a 15 mL tube, placed on ice for 5 min, checked for permeabilization by trypan blue staining (>95%), and then centrifuged in a swinging bucket rotor (1000xg, 4 min, 4°C). Following this, cells were handled using cut tips. The supernatant was removed and samples were resuspended in 1 mL of Cell Wash Buffer (CWB) (10 mM Tris-HCl pH = 8.0, 250 mM Sucrose, 10 mM KCl, 5 mM $MgCl_2$, 1 mM EGTA pH = 8.0, 0.5 mM DTT, 10% Glycerol, 1X protease inhibitor, and 2 µL SUPERase-In RNase inhibitor per 10 mL) and centrifuged (1000xg, 4 min, 4°C) for a total of 2 washes. After the second wash, samples were resuspended in 1 mL total Cell Freeze Buffer (CFB) (50 mM Tris-HCl pH = 8.0, 5 mM $MgCl_2$, 0.5 mM DTT, 40% Glycerol, 1.1 mM EDTA pH = 8.0, and 2 µL SUPERase-In RNase inhibitor per 10 mL), counted by hemocytometer, and then spun down in 1.5 mL tubes in an angled rotor centrifuge (1000xg, 5 min, 4°C). Samples were resuspended in 52 µL of CFB for every 4E6 cells, flash frozen in liquid nitrogen, and then frozen at –80°C until ready to use. For run-on assays, two biotinylated NTP's (UTP and CTP, PerkinElmer, catalog NEL543001EA and NEL542001EA) were used with unbiotinylated ATP and GTP (MilliporeSigma, catalog 11277057001) and 2X ROMM buffer (10 mM Tris-HCl pH = 8.0, 5 mM $MgCl_2$, 1 mM DTT, 300 mM KCl, 40 µM Biotin-11-CTP, 40 µM Biotin-11-UTP, 40 µM ATP, 40 µM GTP, 1% Sarkosyl (MilliporeSigma, catalog L5125) and 1 µL SUPERase-In RNase inhibitor per reaction) was prepared exactly as recommended. 50 µL of pre-heated 2X ROMM buffer was added to 50 µL of cell suspension, pipetted with a cut 200 µL tip 20-25 times quickly, and incubated (with 700 RPM shaking) for 5 min, after which 250 µL of Trizol LS (ThermoFisher Scientific, catalog 10296028) was immediately added, mixed by pipetting, vortexed, and placed on ice. 65 µL chloroform was added to each sample, vortexed, incubated on ice for 3 min, and centrifuged (20,000xg, 8 min, 4°C). Approximately ~150 µL of the aqueous phase was collected in a new tube followed by addition of 1 µL Glycoblue (ThermoFisher Scientific, catalog AM9515) and 2.5X volumes of 100% ethanol, and samples were then vortexed for 5 sec and centrifuged (20,000xg, 20 min, 4°C). The supernatant was removed, washed with 75% ethanol once with gentle inversion and pulse spun. RNA pellets were airdried followed by resuspension in 30 µL of RNAse-free water. RNA was denatured for 30 sec at 65°C, snap cooled on ice, and then fragmented with 7.5 µL of cold 1M NaOH on ice for 10 min. 75 µL of 0.5M Tris-HCl pH = 6.8 was added and mixed by pipetting, and then samples were passed through a calibrated Micro Bio-Spin™ P-30 Gel Columns, Tris Buffer (RNase-free) (Bio-Rad, catalog 7326250) following the manufacturer's instructions. The volume of each sample was brought up to 200 µL with water, and 1 uL glycoblue, 8 µL NaCl, and 500 µL 100%

ethanol was added, vortexed, and then centrifuged (20,000xg, 20 min, 4°C). Ethanol was removed and RNA pellets were stored in –80°C overnight. The next day, 75% ethanol was used to wash the pellet, spun down, and then pellets were resuspended in 6 µL of RNAse-free water. 1 µL of 10 µM VRA3 oligo (See Supplemetary Table 2) was added to the RNA and samples were denatured for 30 sec at 65°C and snap cooled on ice. A buffer containing T4 RNA Ligase 1 (ssRNA ligase) (NEB, catalog M0204L) was added to samples for ligation for 1 hr at 25°C. 10 µL/sample of Dynabeads™ MyOne™ Streptavidin C1 Beads (ThermoFisher Scientific, catalog 65001) were then equilibrated by removing storage buffer by placing on a magnet, washed once in 1 mL bead preparation buffer (0.1 M NaOH and 50 mM NaCl) and twice with 1 mL binding buffer (10 mM Tris-HCl pH = 7.5, 300 mM NaCl, 0.1% Triton X-100, 1 mM EDTA pH = 8.0, and 2 µL SUPERase-In RNase inhibitor per 10 mL). Each wash was done by adding the indicated buffer, flipping the tube on the magnet twice, and then removing the buffer without letting beads dry. Beads were resuspended in 25 µL binding buffer per sample and placed on ice until ready to use. After 3' adapter ligation, 55 µL of binding buffer and then 25 µL of beads was added to each sample and rocked for 20 min at room temperature to bind biotinylated, nascent RNA to the beads. Beads were washed with 500 µL High-Salt Buffer (HSB, 50 mM Tris-HCl pH = 7.5, 0.5% Triton-X-100, 2 M NaCl, 1 mM EDTA pH = 8.0 and 2 µL SUPERase-In RNase inhibitor per 10 mL), then 500 µL Low-Salt Buffer (LSB, 5 mM Tris-HCl pH = 7.5, 0.1% (v/v) Triton X-100, 1 mM EDTA pH = 8.0 and 2 µL SUPERase-In RNase inhibitor per 10 mL), and then resuspended in a buffer containing T4 polynucleotide kinase (NEB, catalog M0201L) and incubated for 30 min at 37°C. Samples then underwent 5' DeCapping using a buffer containing RppH enzyme (NEB, catalog M0356S) for 1 hr at 37°C, which was followed by 5' adapter ligation on-beads (VRA5 oligo (See Supplemetary Table 2) was added to each sample and then RNA-bound beads were denatured as described for 3' adapter ligation) for 1 hr at 25°C. After 5' ligation, the beads were washed with both HSB and LSB, resuspended in 300 µL of Trizol, vortexed, and incubated on ice for 3 min. 60 µL chloroform was added to each sample, vortexed, incubated on ice for 3 min, and then centrifuged (20,000xg, 8 min, 4°C). Approximately ~180 µL of the aqueous phase was collected in a new tube followed by addition of 1 µL Glycoblue and 2.5X volumes of 100% ethanol. Samples were then vortexed and centrifuged (20,000xg, 20 min, 4°C). The supernatant was removed, washed with 75% ethanol once with gentle inversion and pulse spin, and RNA pellets were airdried followed by resuspension in 13.5 µL Reverse Transcriptase (RT) resuspension mix to begin cDNA synthesis, which consisted of 4 µL of 10 µM RP1 oligo (see Supplemetary Table 2) and dNTPs. Samples were denatured at 65°C for 5 min, placed on ice, and resuspended in an RT master-mix consisting of Maxima H Minus RT enzyme (ThermoFisher Scientific, catalog EP0752) and cDNA synthesis was performed on a thermal cycler (50°C for 30 min, 65°C for 15 min, and 85°C for 5 min). 2.5 µL of a 10 µM RP1-X (designated RP1 indexing primer) was added to each cDNA synthesis reaction followed by 78.5 µL of PCR amplification master mix (consisting of Q5® High-Fidelity DNA Polymerase) and PCR was performed with a total of 14 cycles following the conditions (56°C extension) in the original protocol. The samples were purified using 180 µL of KAPA pure beads, and eluted in 15 µL of 10 mM Tris-HCl pH = 8.0. Libraries were quality-checked with Tapestation and Qubit and then sequenced at ~66E6 paired-end reads/sample with 50-bp length using a NextSeq 500 (Illumina) at the UT Southwestern McDermott Center Sequencing Core. If samples had excess adapter dimer contamination, they were subsequently electrophoresed on a 2% agarose gel (80V for 60 min) followed by gel excision in cold room, purification by column (QIAGEN, catalog 28606), and additional TapeStation and Qubit analyses before sequencing.

### Supplementary Tables

#### Supplementary Table 1. Plasmids used in this study.

| Plasmid | Origin |
|---|---|
| pCRIS-PITChv2-C-dTAG-Puro (BRD4) | Addgene 91796 |
| pCRIS-PITChv2-C-TAG-BSD (BRD4) | Addgene 91795 |
| Px330A_sgx _sgPITCh | Universal cutting vector derived from Addgene 58766 and 63670 |
| pCRIS-PITChv2-C-dTAG-Puro (KAP1) | Generated in this study |
| pCRIS-PITChv2-C-dTAG-BSD (KAP1) | Generated in this study |
| Px330A_sgx _KAP1C_sgPITCh | Generated in this study |

**Supplementary Table 2. Primers used in this study.**

| Number | Name | Sequence (5'-3') | Usage |
|---|---|---|---|
| 2691 | KAP1 C-Term FWD Guide | CACCGCATGGGGGCTCCAGCCTCAG | dTAG gRNA cloning |
| 2692 | KAP1 C-Term REV Guide | AAACCTGAGGCTGGAGCCCCCATGC | dTAG gRNA cloning |
| 2810 | KAP1 C-Term gRNA2 PUROBSD FWD | gttccgcgttacatagcatcgtacgcgtacgtgtttggggccctggtgatggcAcctggggtggcggtggctcgggcggtg | dTAG arm cloning |
| 2811 | KAP1 C-Term gRNA2 PURO REV | agcattctagagcatcgtacgcgtacgtgtttggGGCCATGGGGGCTCCAGCCTtcaggcaccgggcttgcgggtcatgcaccaggtg | dTAG arm cloning |
| 2812 | KAP1 C-Term gRNA2 BSD REV | agcattctagagcatcgtacgcgtacgtgtttggGGCCATGGGGGCTCCAGCCTtcagccctcccacacataaccagagggcagcaat | dTAG arm cloning |
| 2548 | pPITChv2_seq fwd | GGGTCATTAGTTCATAGCCC | Sanger seq primer for dTAG arm plasmid |
| 2549 | pPITChv2_seq rev | TATTAGGAAAGGACAGTGGG | Sanger seq primer for dTAG arm plasmid |
| 2550 | pX330S_guide_seq | GCTGGCCTTTTGCTCACATG | Sanger seq primer for dTAG gRNA plasmid |
| 3286 | FOS_RTPCR_FWD | GGGGCAAGGTGGAACAGTTA | RT-qPCR |
| 3287 | FOS_RTPCR_REV | AGTTGGTCTGTCTCCGCTTG | RT-qPCR |
| 3656 | ATF3_RTPCR_FWD | CGCTGGAATCAGTCACTGTCAG | RT-qPCR |
| 3657 | ATF3_RTPCR_REV | CTTGTTTCGGCACTTTGCAGCTG | RT-qPCR |
| 3658 | NR4A1_RTPCR_FWD | GGACAACGCTTCATGCCAGCAT | RT-qPCR |
| 3659 | NR4A1_RTPCR_REV | CCTTGTTAGCCAGGCAGATGTAC | RT-qPCR |
| 3052 | EGR1_ RTPCR_FWD | GGCGAGCAGCCCTACG | RT-qPCR |
| 3053 | EGR1_ RTPCR_REV | GCACCTTCTCGTTGTTCAGAG | RT-qPCR |
| 354 | RPL19_FWD | ATCGATCGCCACATGTATCA | RT-qPCR |
| 355 | RPL19_REV | GCGTGCTTCCTTGGTCTTAG | RT-qPCR |
| 1868 | GAPDH_FWD | GCAAATTCCATGGCACCGT | RT-qPCR |
| 1869 | GAPDH_REV | TCGCCCCACTTGATTTTGG | RT-qPCR |
| 9 | U6_FWD | CTCGCTTCGGCAGCACATATAC | RT-qPCR |
| 10 | U6_REV | GGAACGCTTCACGAATTTGCGTG | RT-qPCR |
| | VRA3 RNA adapter | GAUCGUCGGACUGUAGAACUCUGAAC-/Inverted dT/ | PRO-Seq (Purchased from IDT RNase-free HPLC purified) |
| | VRA5 RNA adapter | CCUUGGCACCCGAGAAUUCCA | See VRA3 usage |
| | RP1 (DNA Oligo) | AATGATACGGCGACCACCGAGATCTACACGTTCAGAGTTCTACAGTCCGA | See VRA3 usage |

**Supplementary Table 3. Antibodies used in this study.**

| Target | Vendor | Catalog Number | Assay (Dilution/time) | Figure |
|---|---|---|---|---|
| KAP1 | Abcam | ab22553 | Western blot (1:2000/1 hr) | Figure 1 and Supplementary Figure 1 |
| HA | BioLegend | 901513 | Western blot (1:2000/Overnight) | Figure 1 and Supplementary Figure 1 |
| c-Fos | Cell Signaling Technologies | 2250 | Western blot (1:1000/Overnight) | Figure 1 and Supplementary Figure 1 |
| ATF3 | Santa Cruz Biotechnologies | sc-81189 | Western blot (1:1000/Overnight) | Figure 1 |
| ELK-1 | Cell Signaling Technologies | 9182 | Western blot (1:500/Overnight) | Supplementary Figure 1 |
| ELK-1 Phos-S383 | Santa Cruz Biotechnologies | sc-8406 | Western blot (1:500/Overnight) | Supplementary Figure 1 |
| Actin Rhodamine | Bio-Rad | 12004163 | Western blot (1:10000/1 hr) | Figure 1 and Supplementary Figure 1 |
| Goat anti-mouse IgG-HRP | Santa Cruz Biotechnologies | sc-2005 | Western blot (1:10000/1 hr) | |
| Donkey anti-rabbit IgG-HRP | Santa Cruz Biotechnologies | sc-2313 | Western blot (1:10000/1 hr) | |
| StarBright Blue 700 Goat Anti-Mouse IgG | Bio-Rad | 12004158 | Western blot (1:10000/1 hr) | |
| HA | Millipore | 05-905 | ChIP (5 μg/ChIP) | Figure 2 |
| RPB3 | Millipore | ABE999 | ChIP (5 μg/ChIP) | Figure 3 |
| SPT5 | Bethyl/Thermo | A300-868A | ChIP (5 μg/ChIP) | Figure 5, Supplementary Figures 6 and 7 |
| CDK9 | Cell Signaling Technologies | 2316 | ChIP (18 μL/ChIP) | Figure 5, Supplementary Figures 6 and 7 |
| SPT5 Phos-T806 | Gift from Robert Fisher lab | | ChIP (7 μL/ChIP) | Supplementary Figure 6 |
| TFIIB | Santa Cruz Biotechnologies | sc-271736 | ChIP (5 μg/ChIP) | Supplementary Figure 7 |
| MED1 | Bethyl | A300-793A | ChIP (5 μg/ChIP) | Supplementary Figure 7 |
| CDK7 | Bethyl | A300-405A | ChIP (5 μg/ChIP) | Figure 5 |
| Ser2P Pol II | Millipore | 04-1571 | ChIP (10 μg/ChIP) | Supplementary Figure 7 |
| Ser5P Pol II | Active Motif | 61085 | ChIP (10 μg/ChIP) | Supplementary Figure 7 |

**Supplementary Table 4. Pearson correlation coefficients for all high-throughput sequencing in this study.**

| Experiment | Number of replicates | Sample** | Pearson's Correlation Coefficient | Figure |
|---|---|---|---|---|
| TT-Seq | 2 | DMSO (8hr) | 0.99 | Supplementary Figure 1 |
| | | dTAG (8 hr) | 1.00 | |
| HA ChIP-Seq | 2 | DMSO-S0 | 0.93 | Figure 2 and Supplementary Figure 2 |
| | | DMSO-S30 | 0.83 | |
| | | DMSO-S180 | 0.84 | |
| | | dTAG-S30 | 0.98 | |
| Pol II ChIP-Seq | 2 | DMSO-S0 | 0.99 | Figure 3 and Supplementary Figure 3 |
| | | DMSO-S15 | 1.00 | |
| | | DMSO-S30 | 0.99 | |
| | | dTAG-S0 | 1.00 | |
| | | dTAG-S15 | 1.00 | |
| | | dTAG-S30 | 0.98 | |
| PRO-Seq | 2 | DMSO-S0 | 0.92 | Figure 4 and Supplementary Figures 4 and 5 |
| | | DMSO-S5 | 0.96 | |
| | | DMSO-S10 | 0.98 | |
| | | dTAG-S0 | 0.95 | |
| | | dTAG-S5 | 0.96 | |
| | | dTAG-S10 | 0.95 | |
| Ser2P Pol II ChIP-Seq | 2 | DMSO-S0 | 1.00 | Supplementary Figure 7 |
| | | DMSO-S30 | 1.00 | |
| | | dTAG-S30 | 1.00 | |

**For TT-Seq, samples were treated with DMSO and dTAG for 8 hr in homeostatic conditions. For all other experiments, cells were treated with serum free media (-S0) or with serum for the indicated time point (for -SX min).

**Supplementary Table 5. Alignment statistics for all high-throughput sequencing in this study.**

### Reads aligning to the human genome (hg38)

| Library | Read pairs examined | Read pair duplicates | Percent duplication |
|---|---|---|---|
| PolII_DMSO_Serum0_Replicate1 | 31139199 | 6294476 | 20.3211 |
| PolII_DMSO_Serum0_Replicate2 | 53092913 | 11960899 | 22.6616 |
| PolII_dTAG_Serum0_Replicate1 | 40657527 | 8301387 | 20.5668 |
| PolII_dTAG_Serum0_Replicate2 | 56973603 | 11277780 | 19.9397 |
| PolII_DMSO_Serum15_Replicate1 | 37015058 | 7463579 | 20.2312 |
| PolII_DMSO_Serum15_Replicate2 | 36368697 | 8062452 | 22.2328 |
| PolII_dTAG_Serum15_Replicate1 | 30761204 | 6251095 | 20.3839 |
| PolII_dTAG_Serum15_Replicate2 | 38423042 | 8041808 | 20.9987 |
| PolII_DMSO_Serum30_Replicate1 | 44639810 | 11855203 | 26.607 |
| PolII_DMSO_Serum30_Replicate2 | 34965948 | 8353523 | 23.9364 |
| PolII_dTAG_Serum30_Replicate1 | 33945856 | 8432169 | 24.8867 |
| PolII_dTAG_Serum30_Replicate2 | 36324858 | 8703529 | 24.0072 |
| HA_DMSO_Serum0_Replicate1 | 40322571 | 9487295 | 23.5932 |
| HA_DMSO_Serum0_Replicate2 | 37981047 | 9096273 | 24.0213 |
| HA_DMSO_Serum30_Replicate1 | 38842070 | 9006254 | 23.2523 |
| HA_DMSO_Serum30_Replicate2 | 36981383 | 8573790 | 23.2457 |
| HA_DMSO_Serum180_Replicate1 | 35130554 | 7742961 | 22.0981 |
| HA_DMSO_Serum180_Replicate2 | 37855181 | 9125958 | 24.1659 |
| HA_dTAG_Serum30_Replicate1 | 33022155 | 7855107 | 23.8401 |
| HA_dTAG_Serum30_Replicate2 | 33463381 | 8099510 | 24.2662 |
| PolII_Ser2P_DMSO_Serum0_Replicate1 | 33022671 | 8739888 | 26.5052 |
| PolII_Ser2P_DMSO_Serum0_Replicate2 | 37910809 | 10874119 | 28.7231 |
| PolII_Ser2P_DMSO_Serum30_Replicate1 | 43140498 | 14149213 | 32.8355 |
| PolII_Ser2P_DMSO_Serum30_Replicate2 | 34302291 | 10480899 | 30.5922 |
| PolII_Ser2P_dTAG_Serum30_Replicate1 | 42211318 | 15464820 | 36.6681 |
| PolII_Ser2P_dTAG_Serum30_Replicate2 | 39556026 | 13596583 | 34.4069 |
| PolII_Ser5P_DMSO_Serum0 | 38517462 | 9335665 | 24.3913 |
| PolII_Ser5P_DMSO_Serum30 | 37912478 | 7384822 | 19.5966 |
| PolII_Ser5P_dTAG_Serum30 | 39801598 | 7195715 | 18.1929 |
| SPT5_DMSO_Serum0 | 24784351 | 6514407 | 26.367 |
| SPT5_DMSO_Serum5 | 25693981 | 4651674 | 18.2227 |
| SPT5_dTAG_Serum5 | 25874588 | 5152945 | 20.0323 |
| SPT5_DMSO_Serum30 | 27387149 | 5717553 | 20.9875 |
| SPT5_dTAG_Serum30 | 26033518 | 6025396 | 23.2462 |
| pSPT5_Thr806P_DMSO_Serum0 | 29791089 | 11146540 | 37.4839 |
| pSPT5_Thr806P_DMSO_Serum5 | 25326385 | 8855041 | 35.0331 |
| pSPT5_Thr806P_dTAG_Serum5 | 25551680 | 6274354 | 24.6525 |
| CDK9_DMSO_Serum0 | 27375650 | 18760106 | 68.5339 |
| CDK9_DMSO_Serum5 | 27834349 | 14433264 | 51.886 |
| CDK9_dTAG_Serum5 | 25574821 | 16180859 | 63.2805 |
| CDK9_DMSO_Serum30 | 22487188 | 9311695 | 41.4603 |
| CDK9_dTAG_Serum30 | 25161269 | 12385526 | 49.2601 |
| CDK7_DMSO_Serum0 | 39339209 | 14340605 | 36.5163 |
| CDK7_DMSO_Serum30 | 34799419 | 14368796 | 41.3568 |
| CDK7_dTAG_Serum30 | 38087252 | 15085153 | 39.6802 |
| MED1_DMSO_Serum0 | 40415656 | 27890930 | 69.0142 |
| MED1_DMSO_Serum30 | 35591514 | 24323052 | 68.3405 |

| | | | |
|---|---|---|---|
| MED1_dTAG_Serum30 | 36008872 | 24138715 | 67.0311 |
| TFIIB_DMSO_Serum0 | 20164961 | 15605622 | 77.3531 |
| TFIIB_DMSO_Serum30 | 21264825 | 12008670 | 56.4894 |
| TFIIB_dTAG_Serum30 | 23295485 | 15770857 | 67.6928 |

### Spike-in reads aligning to the Drosophila genome (Dm6)

| ChIP-Seq library | Read pairs examined | Read pair duplicates | Percent duplication | Spike-in Scale Factors applied to human files |
|---|---|---|---|---|
| HA_DMSO_Serum0_R1 | 36323 | 7994 | 39.084 | 1.251 |
| HA_DMSO_Serum0_R2 | 40994 | 9010 | 38.5216 | 1.251 |
| HA_DMSO_Serum30_R1 | 36214 | 7482 | 38.6249 | 1.251 |
| HA_DMSO_Serum30_R2 | 21170 | 4249 | 42.2422 | 1.3335 |
| HA_DMSO_Serum180_R1 | 15691 | 3128 | 43.0812 | 1.3643 |
| HA_DMSO_Serum180_R2 | 18918 | 4022 | 43.4092 | 1.3335 |
| HA_dTAG_Serum30_R1 | 147108 | 32323 | 26.2654 | 0.4548 |
| HA_dTAG_Serum30_R2 | 167175 | 37739 | 26.6953 | 0.6821 |
| SPT5_DMSO_Serum0 | 79289 | 21707 | 32.2842 | 0.7579 |
| SPT5_DMSO_Serum5 | 29941 | 5183 | 32.0865 | 1.1487 |
| SPT5_dTAG_Serum5 | 27491 | 5339 | 34.4706 | 1.1789 |
| SPT5_DMSO_Serum30 | 40127 | 8249 | 33.569 | 1.1487 |
| SPT5_dTAG_Serum30 | 43116 | 9935 | 34.0166 | 1 |
| pSPT5_Thr806P_DMSO_Serum0 | 308173 | 126015 | 41.9428 | 0.7937 |
| pSPT5_Thr806P_DMSO_Serum5 | 99482 | 36701 | 40.1875 | 1.2114 |
| pSPT5_Thr806P_dTAG_Serum5 | 95869 | 24542 | 30.8381 | 1.1447 |
| CDK9_DMSO_Serum0 | 452080 | 333921 | 73.5347 | 1.0073 |
| CDK9_DMSO_Serum5 | 245142 | 135489 | 55.8562 | 1 |
| CDK9_dTAG_Serum5 | 233371 | 155879 | 66.6648 | 1.2941 |
| CDK9_DMSO_Serum30 | 210504 | 96449 | 46.8914 | 0.9441 |
| CDK9_dTAG_Serum30 | 306263 | 165895 | 54.6788 | 0.7841 |
| CDK7_DMSO_Serum0 | 112910 | 44695 | 44.4832 | 1 |
| CDK7_DMSO_Serum30 | 130406 | 56403 | 47.1684 | 1 |
| CDK7_dTAG_Serum30 | 156589 | 65781 | 46.2314 | 1 |
| MED1_DMSO_Serum0 | 561891 | 404285 | 72.9469 | 0.9103 |
| MED1_DMSO_Serum30 | 505410 | 358961 | 71.6832 | 0.951 |
| MED1_dTAG_Serum30 | 450515 | 316463 | 70.6458 | 1 |
| TFIIB_DMSO_Serum0 | 189012 | 148324 | 77.4249 | 1 |
| TFIIB_DMSO_Serum30 | 186992 | 116588 | 62.159 | 0.9086 |
| TFIIB_dTAG_Serum30 | 115949 | 82997 | 70.4418 | 1.1447 |
